# Speech-in-noise detection is related to auditory working memory precision for frequency

**DOI:** 10.1101/2020.01.22.915165

**Authors:** Meher Lad, Emma Holmes, Agatha Chu, Timothy D Griffiths

## Abstract

Speech-in-noise (SiN) perception is a critical aspect of natural listening, deficits in which are a major contributor to the hearing handicap in cochlear hearing loss. Studies suggest that SiN perception correlates with cognitive skills, particularly phonological working memory: the ability to hold and manipulate phonemes or words in mind. We consider here the idea that SiN perception is linked to a more general ability to hold sound objects in mind, auditory working memory, irrespective of whether the objects are speech sounds. This process might help combine foreground elements, like speech, over seconds to aid their separation from the background of an auditory scene.

We investigated the relationship between auditory working memory precision and SiN thresholds in listeners with normal hearing. We used a novel paradigm that tests auditory working memory for non-speech sounds that vary in frequency and amplitude modulation (AM) rate. The paradigm yields measures of precision in frequency and AM domains, based on the distribution of participants’ estimates of the target. Across participants, frequency precision correlated significantly with SiN thresholds. Frequency precision also correlated with the number of years of musical training. Measures of phonological working memory did not correlate with SiN detection ability.

Our results demonstrate a specific relationship between working memory for frequency and SiN. We suggest that working memory for frequency facilitates the identification and tracking of foreground objects like speech during natural listening. Working memory performance for frequency also correlated with years of musical instrument experience suggesting that the former is potentially modifiable.

## Introduction

Speech-in-noise (SiN) perception is the ability to identify spoken words when background noise is present. Deficits in SiN are one of the most common problems in patients with cochlear hearing loss, but there has been increasing interest in cognitive abilities that determine SiN perception^1^. Akeroyd^2^ summarised studies describing the relationship of ccognitive measures to speech-in-noise performance. Phonological working measures such as the reading span and digit span were found to have an effect on SiN detection after accounting for hearing loss. However, other studies have suggested that phonological working memory only comes into play in older participants or when a participant has high-frequency hearing loss^3^.

We consider here the idea that more fundamental forms of working memory that apply to all sounds, including speech, are relevant to SiN perception. From first principles, the ability to hold in mind sound features that are characteristic of particular sources, including voices, might aid SiN perception by allowing sequential outputs from a particular source to be grouped. Previous work has shown that fundamental auditory grouping processes involved in separating non-speech figures from an acoustic background (‘figure-ground perception’) explains a sizable portion of individual differences in SiN ability^4^; however, not all of the variance in SiN was accounted for by these processes. The current work investigates another aspect of auditory cognition that might help to account for individual differences in SiN ability: auditory working memory for frequency and amplitude modulation. These working memory and figure-ground perception measures both allow robust psychophysical characterisation with the potential to predict SiN in a way that is independent of language and education. Previous work^5–7^ has shown that tasks measuring the ability to segregate sound based on frequency characteristics of complex sounds can be related to individual SiN perception ability. However, this effect has only been found in aged participants or those with hearing loss.

In this work, we measured the precision of working memory for the frequency and modulation rate of amplitude-modulated narrowband sounds. The task (see Methods section for more details) tests a participant’s ability to match a pure tone to one that had been presented a few seconds earlier. The matching process is repeated over 100 trials and the errors between the matched sound and the original sound are used to calculate the overall precision of that participant as the inverse of the standard deviation of the errors. This approach has been developed from models of working memory initially developed in the visual domain, in which working memory is treated as a finite resource that can be distributed over multiple objects with a degree of precision determined by the number of objects^7^. Previous work^9^ in the auditory system also supports the use of a resource model applied to frequency and amplitude modulation rate in this way. We were interested to investigate potential differences between frequency as a property of sources (like voices) and modulation rate as a property of events (like words), where the linking of source characteristics might better predict SiN ability.

We recruited 44 participants with normal hearing (further participant details in the Methods section), defined by a strict criterion of audiometric thresholds ≤ 15 dB HL in either ear at octave frequencies between 0.25 and 8 kHz. We measured their thresholds for reporting sentences from a closed-set speech corpus spoken by a male talker in the presence of 16-talker babble. Given that previous studies have reported relationships between SiN and phonological working memory^2^, audiometric thresholds^4, 10^, and age^11^, we examined these relationships in addition to working memory precision. In addition, we asked participants about their musical training history, which we predicted might relate to SiN or to working memory ability. The idea that musicians have better SiN detection has been controversial^12–14^, with recent evidence suggesting this is not related to SiN detection directly^15^. However, the additional cognitive demands of working memory tasks might be more relevant to musical listening^16^.

We found that SiN thresholds correlate with the precision of working memory for frequency, but not with the precision of WM for AM or phonological WM. The precision of WM for frequency also correlates with years of musical training, raising the idea that this could be a trainable skill.

## Results

44 participants without any neurological or psychiatric history were tested. The audiometric thresholds ranged from -5.8 to 14.2 dB HL with a mean value of 7.1 dB HL and standard deviation of 5.0 dB HL.

### Relationship between SiN and hearing thresholds

We tested the relationship between audiometric thresholds at 4–8 kHz and SiN thresholds given a previous demonstration of a significant relationship between these variables^4^. We did not find a significant correlation between audiometric thresholds in this range and SiN perception ability (r = 0.31, p = 0.176).

### Relationship between SiN and auditory WM

Frequency precision values were not normally distributed, so a Spearman’s rank correlation was used for these analyses. Figure 1 shows the relationships between SiN performance and the auditory WM precision measures. Bootstrapping based on 1000 samples was performed to estimate the 95% confidence intervals for the correlation coefficients for working memory metric correlations for frequency and AM with SiN thresholds. SiN thresholds correlated significantly with the precision measure of auditory WM for frequency (ρ = -0.36 (CI = -0.622 to -0.061), p = 0.016) (Fig 1A). This remained statistically significant after Bonferroni-Holm correction. AM precision (r = -0.22 (CI = -0.482 to 0.129), p = 0.154) (Fig 1B) did not correlate significantly with SiN performance, although there was no significant difference in the correlation coefficients between the frequency and AM metrics and SiN performance (*t* = 01.589, p = 0.119).

**Fig 1.**
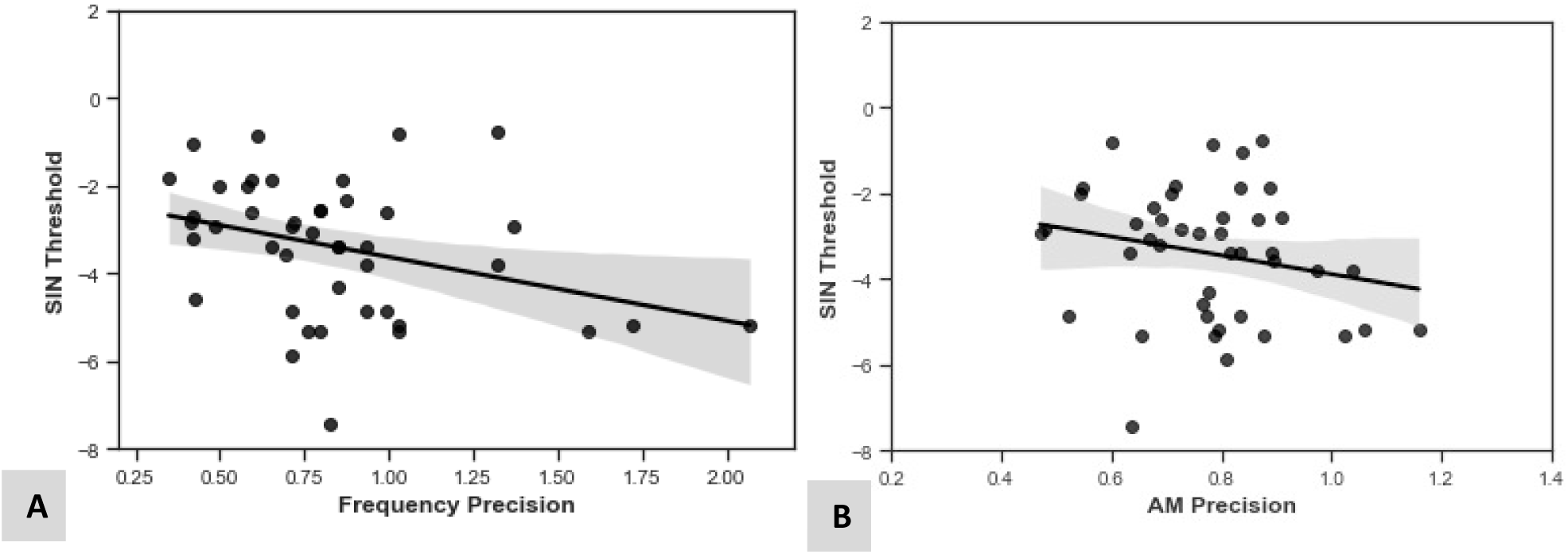
Relationships between SiN performance and the auditory WM measures. Black dots show the results from individual participants and the black lines show the least squares lines of best fit. The grey shaded areas shows the standard error. Confidence intervals for the correlation coefficients were calculated by performing bootstrapping with 1000 samples. (A) Better SiN ability (lower thresholds) is significantly correlated with higher frequency WM precision (ρ = -0.36 (CI = -0.622 to -0.061), p = 0.016). (B) There was no significant association between SiN ability and AM WM precision (r = -0.22 (CI = -0.482 to 0.129), p = 0.154).

We used multiple regression to determine the contributions of peripheral and central auditory measures to SiN as done previously in a larger cohort^4^. A model including hearing thresholds and auditory WM precision for frequency (r = -0.35, r^2^ change = 0.005, p-value change = 0.065) showed only a marginal increase in variance explained than with the WM measure alone.

### Relationship between SiN and phonological working memory measures

We used conventional forward and backward digit span measures of phonological working memory. We tested the relationship between scores on these tests and SiN thresholds. The results of these correlations— and of correlations between SiN and reading ability and reasoning scores—are displayed in Table 1. Crucially, the forward (r = -0.12, p = 0.490) and backward (r = -0.32, p = 0.069) digit span measures did not correlate significantly with SiN thresholds.

**Table 1.** R-values and P-values for correlations between speech-in-noise (SIN) detection ability, auditory working memory (AWM) and Neuropsychometric scores. Reading ability was measured using the Weschler Test of Adult Reading. Non-verbal reasoning was measured using the Block Design subsection of the Weschler Adult Intelligence Scale 3. Delayed verbal recall was measured using the List Learning A set from the Weschler Adult Intelligence Scale 3. Asterisks indicate statistically significant correlations after correction for multiple comparisons using the Bonferroni-Holm method.

|  | **SIN DETECTION** |  | **FREQUENCY AWM** |  | **AM AWM** |  |
| --- | --- | --- | --- | --- | --- | --- |
| **FORWARD DIGIT SPAN** | -0.12 | -0.490 | 0.36 | 0.027 | 0.14 | 0.420 |
| **BACKWARD DIGIT SPAN** | -0.32 | 0.069 | 0.50 | 0.002* | 0.27 | 0.110 |
| **READING ABILITY** | -0.50 | -0.003* | 0.29 | 0.082 | 0.22 | 0.191 |
| **NON-VERBAL REASONING** | 0.10 | 0.590 | 0.31 | 0.059 | 0.19 | 0.260 |
| **DELAYED VERBAL RECALL** | -0.04 | -0.810 | -0.10 | 0.560 | 0.10 | 0.560 |

We also examined the relationships between these measures and auditory working memory (Fig 2). Only backward (r = 0.50, p = 0.002; Fig 2B) digit span correlated with the precision of WM for frequency after correction. Neither correlated with the precision of WM for AM (forward span: r = 0.14, p = 0.415; backward span: r = 0.27, p = 0.109).

**Fig 2.**
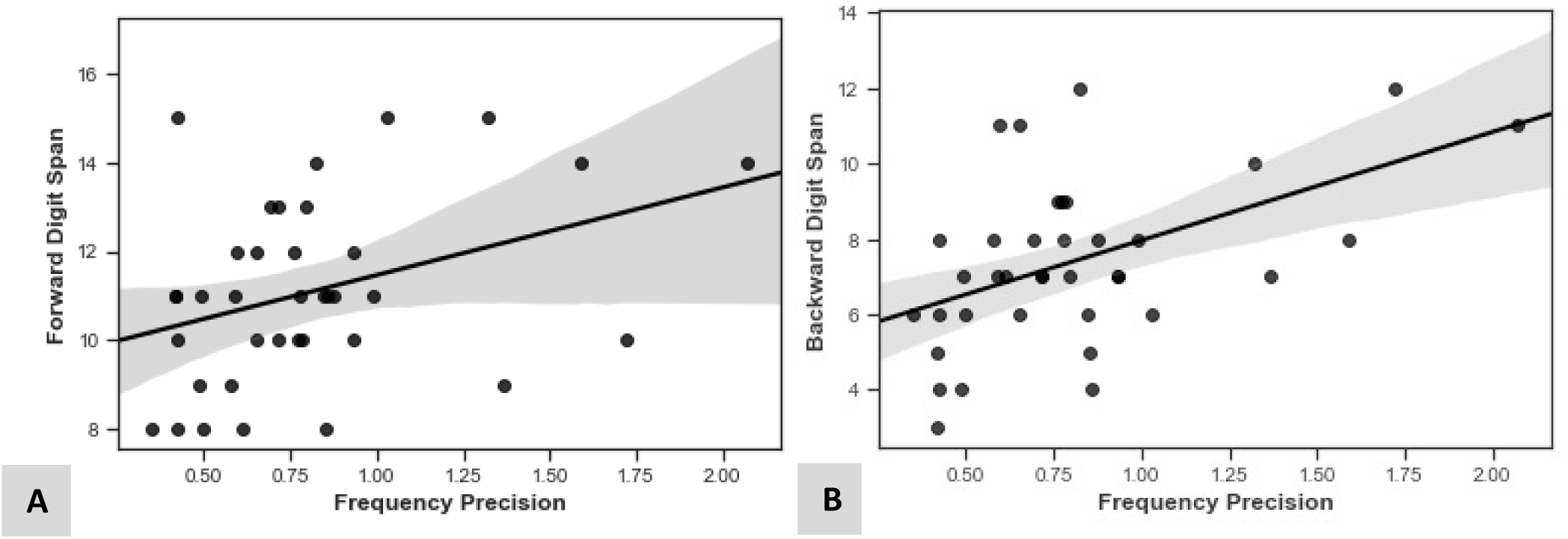
Relationships between precision of WM for frequency and phonological WM. (A) Forward digit span (r = 0.36, p = 0.027) and (B) backward digit span (r = 0.50, p = 0.002).

### Relationships with age and years of musical experience

We tested the effect of age on the two auditory WM precision metrics: frequency and AM. Age did not satisfy the Shapiro-Wilk test for normality, so we used Spearman’s rank correlations for this variable. Age correlated positively with hearing thresholds (ρ = 0.60, p < 0.001), but it did not correlate significantly with SiN thresholds (ρ = 0.01, p = 0.974), frequency WM precision (ρ = -0.20, p = 0.180) or AM WM precision (ρ = -0.06, p = 0.701).

We documented musical training for each participant as the self-reported number of years (years from starting to stopping playing an instrument or to current date if they were still playing) a participant has played a musical instrument. Participants who did not play an instrument were coded as zero. The values for this variable were not normally distributed, so non-parametric tests were used. Years of musical training showed a significant correlation with frequency WM precision (ρ = 0.57, p < 0.001; Fig 3) but not AM WM precision (ρ = 0.22, p = 0.192). There was no significant correlation between SiN performance and years of musical training (r = 0.01, p = 0.960).

**Fig 3.**
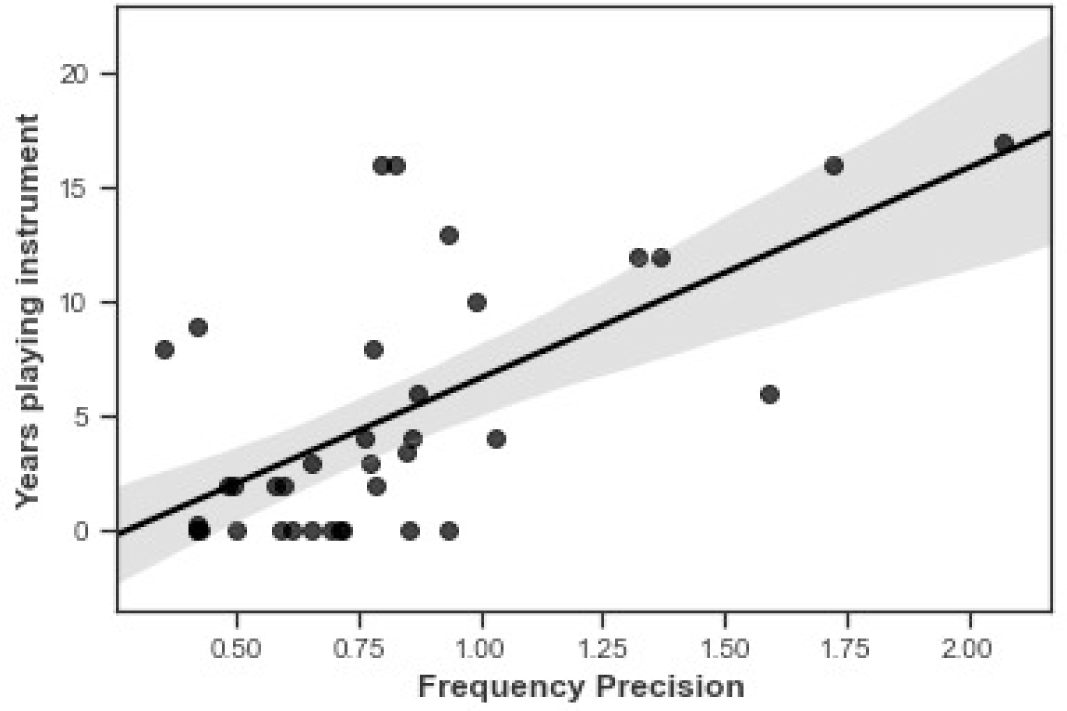
Positive correlation between the number of years a participant has played a musical instrument and frequency WM precision (ρ = 0.57, p < 0.001).

## Discussion

We provide evidence that auditory WM for frequency is an important factor in SiN perception. In this study, WM precision for frequency correlated significantly with thresholds for reporting sentences in multi-talker babble. The association with frequency precision highlights the importance of frequency cues in the perception of target speech from background noise. In contrast to some previous studies, peripheral audiometric hearing measures^4^ and phonological working memory^11^ measures were not significantly associated with SiN ^4,2,1^. In addition, performance in the WM task for frequency precision correlated with years of musical training, potentially implicating musical ability as a modifiable factor for auditory WM.

### SiN and hearing loss

Previous studies have identified that hearing loss, as measured by the pure-tone audiogram, is a strong correlate of SiN ability among people with hearing loss^8^, and may also explain some of the variability in SiN performance among people who have audiometric thresholds at clinically measured frequencies (0.25–8 kHz) in the normal hearing range^4^. There is also evidence that extended high-frequency hearing (8 to 20 kHz) can enhance SiN perception^18^. Although pure-tone audiometric thresholds may be used synonymously with ‘hearing loss’, this test measures tone-sensitivity in quiet, which is different from SiN. For example, even when tone-sensitivity thresholds are corrected by a hearing aid, up to 15% of people report difficulty when communicating in noisy environments^11^. Given that neuroimaging studies demonstrate that SiN perception evokes cortical activity, peripheral damage may be anatomically distinct from some SiN deficits^12^.

Here, we were interested in SiN in people with normal hearing and we utilised a stringent criterion to exclude participants with mild tone sensitivity impairment in the high-frequency range. We did not measure audiometric thresholds above 8 kHz, which may have some effect on SiN^13,14^ because these thresholds are not measured routinely in clinical practice. Although, there was a relationship between high-frequency (4–8 kHz) audiometric thresholds and SiN in the full sample, consistent with the results of Holmes and Griffiths^4^, we did not find a significant correlation between audiometric hearing thresholds and SiN performance after excluding participants with audiometric thresholds > 15 dB HL. This may be because our participants were younger than in that previous experiment. One previous experiment that used a similar age range found no correlation^17^, and one previous experiment that used a similar threshold criterion found no correlation^18^. In addition, our sample size was smaller than the experiment reported by Holmes and Griffiths^4^ and therefore the absence of a correlation may be explained by lower power for detecting significant correlations, particularly after excluding participants.

### SiN performance and auditory working memory

The novel auditory WM tasks used here tested specific aspects of WM and did not include any linguistic aspects, which could affect performance. We found that frequency WM precision correlated significantly with SiN thresholds, but AM WM precision did not. However, the difference in correlations between frequency precision and SiN, and AM precision and SiN was not statistically significant. Our findings indicate that people who are best able to hold in mind frequency over time, arguably a source property (as opposed to the temporal envelope related to events), have an advantage in SiN perception.

There is little information on how higher precision for frequency may aid SiN perception, but clues can be drawn from some previous studies. Previous studies using a small number of competing talkers have shown that talkers who are separated from maskers in fundamental frequency are more intelligible than talkers who are closer in pitch to maskers^26,27^. Thus, at a broad scale, it seems plausible that the ability to hold frequency information in memory contributes to speech intelligibility in noise. The frequency of sounds in an auditory scene may also help group them into an auditory object and the precision at which one may be able to do this over time could aid the sequential segregation needed for successful SiN perception^22^. We may have expected AM WM to correlate with SiN. However, it is worth noting here that the SiN masker we used was 16-talker babble and therefore contained less prominent AM than a single-talker masker. AM WM may potentially come into play when a fluctuating masker means that temporal glimpses of the target are more prevalent.

We designed our task to specifically test auditory WM and attempted to minimise the influence of perception during the matching phase of the task. For example, both the frequency and AM rate could only be altered by a step increment of either +2% or -2% in the matching phase of a trial, which is above the discrimination thresholds for frequency and AM rate^23^. However, we did not formally measure the difference limens for each participant. Further work is needed to study how frequency discrimination is related to working memory precision with our task. This may also be relevant for participants with tone deafness who may have lower perceptual thresholds than the normal population and may therefore have been impaired in our task.

We suspect that the WM measures used here tap into different processes than the fundamental grouping processes measured by Holmes and Griffiths^4^. Those processes are carried out over hundreds of milliseconds as opposed to the working memory precision examined here that examines a process carried out over seconds. It will be interesting in future studies to examine the extent to which the figure ground task and auditory working memory tasks explain separate variance in SiN.

### SiN performance and cognitive measures

Several studies have shown that that cognitive function is correlated with SiN ability^19,20^. These studies have found relevance in measures of fluid intelligence, processing speed, and working memory batteries. A large UK Biobank looked at correlations between the Triple-Digits Task (TDT), as a measure of SiN, and the above measures in half a million participants in midlife. They found a significant relationship with all cognitive measures independent of tone-sensitivity, as age increases from 40 to 70 years. A large meta-analysis^29^ looking at the link between cognitive function and SiN perception found that when all cognitive domains and SiN perception tasks were collapsed across all studies, they correlated significantly (r = 0.31). Inhibitory control, WM, episodic memory and processing speed were all deemed to be important correlates of SiN.

From first principles, there are multiple cognitive factors that could plausibly affect SiN perception. A listener must attend to the correct auditory stream, then track and remember this over time. There may also be top-down effects of language which may help anticipate and resolve any ambiguities that are encountered. However, the extent to which individual differences in these cognitive abilities relate to individual differences in speech-in-noise perception among young and healthy participants is equivocal. A review by Akeroyd^2^ suggested that no single test produces reliable correlations, although there is perhaps a more reliable link with phonological WM than with other measures.

We found that a verbal reading test correlated significantly with SiN performance. However, non-verbal measures, such as block design, did not correlate and neither did measures of long-term memory, such as list learning (see Table 1). This is consistent with evidence suggesting that linguistic experience can have an effect on speech discrimination due to advantages in lexical access^21^. Previous work has shown a wide variation when measures of crystallised intelligence have been used. The overall pooled association to SiN perception tasks that have been used in the literature was 0.19, however, this increased when SiN tasks which required a listener to correctly identify sentences, as opposed to words, was used. Within this meta-analysis, one study found a much stronger correlation than ours of 0.70 when the scores from the vocabulary section of participants from the Weschler Adult Intelligence Scale was correlated with performance on a SiN perception task using a background of multi-talker babble with 20 speakers. However, some other studies have not found a significant relationship with tests of intelligence^27,31^.

Considering phonological working memory tests, we found no correlation between SiN performance and forward or backward digit span although this has been identified as a significant correlate of SiN performance in large studies with participants who are older and/or have hearing loss^22^. Researchers have used a variety of tests, including the reading span, forward and backward digit span. Some but not all studies have found significant correlations between the reading span and SiN performance^18,19^. There have also been studies finding a positive association with the digit span and SiN measures^25^. Neither the forward or backward digit span correlated with SiN performance in our study but the strength of association between the backward digit span (r = 0.32) was similar to that in meta-analyses (r = 0.31)^29^. One of the reasons for this may be the smaller sample size of our participants compared to the studies discussed above. The fine-grained measure of WM precision may also be more sensitive to detecting trends that the traditional phonological measures. Additionally, the mean level of hearing loss in these studies was around 40 dB HL whereas it was 7.1 dB HL in our study, which could have influenced this finding as suggested by Füllgrabe and Rosen^3^.

### Role of musical instrument experience in working memory and SiN performance

There is a large body of literature which has identified that musical instrument experience modulates cognitive ability. Playing a musical instrument has been associated with better global cognitive scores as one gets older^26^, whereas this relationship is absent in musicians not actively playing an instrument. A twin study^36^ has also shown that the twin that played an instrument were less likely to develop dementia later in life. Musical training involves the simultaneous use of auditory, motor and visuospatial abilities and these use of these abilities in tandem over time may have general protective effects on the brain.

Although our auditory working memory task for frequency precision requires the maintenance and comparison of tones in the musical range, over seconds, it does not include other aspects of music including melody or rhythm. However, playing music does require a sustained period of working memory for sound which is reflected in our task. Therefore, performance in our task may have been directly related to this. Further work is necessary to clarify whether the nature of musical expertise, the intensity of training and/or the type of instrument that is learned affects frequency working memory precision. It is also possible that the relationship may be simply driven by a tendency for people with better sensitivity for frequency to persist in musical activities.

Musical training has also been linked specifically with better SiN performance and frequency discrimination although this is controversial^13,26,27,15^. A study by Parbery-Clark *et al*.^37^ found that years of musical experience correlated with phonological working memory and frequency discrimination, as well as SiN. It is worth noting that their frequency discrimination task may contain elements of working memory. They used an adaptive paradigm to obtain a threshold for identifying the ‘higher’ sound out of two tones played in a sequence. Although this is described as a perceptual task, this task requires participants to maintain both tones over a few seconds, and compare them in memory (however, explicit details about timing are not mentioned in the study). It is plausible that the expertise gained from musical training in analysing and attending to auditory streams on a background of different music is transferable into more general and non-musical domains of SiN perception and cognition. To this end, further work is needed to establish whether expertise in attending to auditory streams of music in a background of other music (i.e. playing an instrument in an orchestra) adds an advantage above doing so without background music (i.e. soloists). This is a limitation of our study as the participants did not provide further details about the nature of their previous experience with a musical instrument (e.g. amateur vs. professional, self-taught vs. formal musical training). There is also other work that has not found a direct link between musical ability and SiN^37^.

## Conclusions

We present a novel auditory working memory paradigm that acts as a sensitive correlate of SiN performance. We show that the precision of working memory for frequency is significantly related to SiN performance in people with normal audiometric thresholds at standard clinical frequencies. Frequency precision, but not AM precision, correlates with phonological working memory and the former is a better predictor of SiN ability. Additionally, we found that years of musical training was a strong predictor of the precision of working memory for frequency.

## Methods

### Participants

44 participants (19 male, age range 18–53) were recruited from the Institute of Neuroscience, Newcastle University participant database. Participants had a mean age of 30 years with a standard deviation of 9 years. None of the participants had a significant neurological or psychiatric diagnosis. Participants were compensated for their time at the rate of £10 an hour.

Informed written consent was obtained for all participants. This study was approved by the Newcastle University Research Ethics Committee and all research was performed in accordance with local university guidelines and regulations.

### Audiometry

All participants underwent a pure-tone audiogram to quantify any clinically relevant hearing loss. We measured thresholds at octave frequencies between 0.25 and 8 kHz (inclusive) in accordance with BD EN ISO 8253-I (British Society of Audiology, 2004). Participants had subclinical measurements for high frequency averages (4–8 kHz) which were used in subsequent analyses.

### Experimental Methods

44 participants performed all of the auditory tasks. A subset, 33 participants performed neuropsychological tests as well.

Neuropsychological tests included the Weschler Test of Adult Reading (WTAR); List Learning from the BIRT Memory and Information Processing Battery; Forward, Reverse Digit Span and Block Design from the Weschler Adult Intelligence Scale 3 Test Battery.

Auditory tasks were performed in a sound-proof room in the Auditory Laboratory, Institute of Neuroscience, Newcastle University. These consisted of a SiN task and novel auditory WM tasks. Stimuli were presented through circumaural headphones (Sennheiser HD 201) at 70 dB A. All tasks were programmed and developed using the Psychtoolbox package in MATLAB and the PsychoPy module in Python^39^.

The SiN task was completed in two separate blocks, which each contained 2 interleaved runs. Each run contained different target sentences. Participants heard target sentences simultaneously with 16-talker babble. The babble began 500ms before the sentence began and ended 500ms after it ended. A different segment of babble was presented on each trial. Target sentences had the form <name> <verb> <number> <adjective> <noun> (e.g. “Cathy bought four large chairs” and “Peter got two large desks”) and participants were asked to report the five words from the sentence by clicking words (10 options for each word) on a screen in any order. A 1-up, 1-down adaptive paradigm was implemented with a starting signal-to-noise ratio of 0 dB. All 5 words had to be correctly identified for the signal-to-noise ratio to decrease. Step sizes decreased from 2 dB to 0.5 dB after 3 reversals and each run terminated after 10 reversals. The last 6 reversals were used to calculate the threshold for each run, and the thresholds for the 4 runs were averaged to calculate the SiN measure for each participant.

The auditory WM task tests working memory precision for pure tones that differ in their frequency (400– 1000 Hz) and amplitude-modulation rate (4–16 Hz), each containing 100 trials. On each trial, participants listened to two sounds with an inter-stimulus interval of 1–4 seconds. Their task was to match the frequency or AM of the second sound to the first. Participants only heard the stimulus pair once, at the beginning of the trial. They were instructed to press the left and right arrow keys on a keyboard to alter the percept (by varying the frequency or AM rate) of the sound until they perceived that the second sound was the same as the first. Every time an adjustment was made with the keyboard, the participant would hear the ‘adjusted’ sound until they ‘accepted’ the sound as a match for that trial. For example, for the frequency-matching task, the left arrow key would decrease the frequency and the right would increase the frequency, leading to a different pitch percept. Changes in frequency took place in steps of 2% which is above the perceptual threshold for sound discrimination. Changes in AM rate took place in steps of 1 Hz, which in pilot studies was easily discriminated. In the task in which both frequency and AM rate could change, left and right arrow keys altered the frequency, whereas up and down arrow keys altered the AM rate. Performance on each trial was evaluated as the error (i.e., difference) between the parameter of the first sound and that of the perceptually matched second sound. The precision was calculated across trials as the inverse of the standard deviation in error. According to published evidence^8,9^, this better reflects working memory processes according to newer models. The participants performed each task in 2 blocks, and the blocks from the three tasks were pseudorandomly interleaved. The order of these blocks was counterbalanced across participants, so that the order was matched as closely as possible between tasks.

### Data Analysis

For the pure-tone audiogram, a 2-frequency average at 4–8 kHz across the left and right ears was used for the analyses. This was based on previous evidence from our group showing subclinical hearing thresholds in this range correlating significantly with speech-in-noise perception ability using the same task used in the current study^4^, and because variability in the current group of participants was small for low-frequency thresholds. For neuropsychological tests, we entered absolute scores into the analyses. For the SiN task, we took the mean of thresholds in the 4 runs.

We used Pearson’s correlation coefficients to assess relationships between the variables of interest, except where the assumptions were violated where Spearman’s rank correlation was used. This is indicated in the results section. Bootstrapping based on 1000 samples was performed to estimate the 95% confidence intervals for the correlation coefficients for working memory metric correlations for frequency and AM with SiN thresholds. The Bonferroni-Holm method was used to adjust Family-Wise Error Rate. The Steiger method was used to test a difference in correlation coefficients using the *corrstats* module in Python. All other analyses were performed in IBM SPSS Statistics Version 24.

## Acknowledgements

We would like to thank all the volunteers that took part in the study. ML is funded by the Guarantors of Brain Entry Fellowship. EH is funded by Action on Hearing Loss Pauline Ashley Fellowship scheme. AC was funded by a Newcastle University Vacation Scholarship. TDG is funded by a Senior Fellowship grant from the Wellcome Trust.

All authors have no conflicts of interests to declare and all data is available at request.

